# Proper reference selection and re-referencing to mitigate bias in single pulse electrical stimulation data

**DOI:** 10.1101/2024.10.21.619449

**Authors:** Harvey Huang, Joshua A. Adkinson, Michael A. Jensen, Mohammed Hasen, Isabel A. Danstrom, Kelly R. Bijanki, Nicholas M. Gregg, Kai J. Miller, Sameer A. Sheth, Dora Hermes, Eleonora Bartoli

## Abstract

Single pulse electrical stimulation experiments produce pulse-evoked potentials used to infer brain connectivity. The choice of recording reference for intracranial electrodes remains non-standardized and can significantly impact data interpretation. When the reference electrode is affected by stimulation or evoked brain activity, it can contaminate the pulse-evoked potentials recorded at all other electrodes and influence interpretation of findings. We highlight this specific issue in intracranial EEG datasets from two subjects recorded at separate institutions. We present several intuitive metrics to detect the presence of reference contamination and offer practical guidance on different mitigation strategies. Either switching the reference electrode or re-referencing to an adjusted common average effectively mitigated the reference contamination issue, as evidenced by increased variability in pulse-evoked potentials across the brain. Overall, we demonstrate the importance of clear quality checks and preprocessing steps that should be performed before analysis of single pulse electrical stimulation data.

**Highlights:**

- A reference electrode close to active tissue can contaminate intracranial EEG signals
- Interpretation of pulse-evoked potentials can be biased by reference contamination
- Low response variability between channels is indicative of reference contamination
- Reference contamination can be resolved through data recollection or re-referencing

## Introduction

Single pulse electrical stimulation (SPES) experiments assess functional effective connectivity in human intracranial recordings by testing for causal influence between brain regions (Kundu et al., 2020; Matsumoto et al., 2004). The pulse-evoked potentials (PEPs) measured at distant electrode locations from the stimulation site are quantified (e.g., by amplitude, latency, or overall shape) and interpreted as a proxy of directed causal connectivity for each pair of stimulated-recorded locations (Crocker et al., 2021; Huang et al., 2023; Miller et al., 2021; Schmid et al., 2024). When stimulation and recording are done via subdural cortical electrodes, these are more commonly referred to as “cortico-cortical evoked potentials”.

Methods for preprocessing SPES data are inconsistent across researchers, and can significantly impact data interpretation (Levinson et al., 2024). For example, a carefully chosen hardware reference and *post hoc* re-referencing scheme are needed to highlight neural features of interest in the data (Huang et al., 2024; Li et al., 2018; Mercier et al., 2022). Re-referencing ensures comparability of findings across datasets, but it may also alter the original recorded signals and change the average PEP waveform. When the reference electrode is positioned close to either the stimulated electrodes or a brain area with a robust evoked response, the raw data can be contaminated by artifacts or signals. Re-referencing can remove some of these unwanted evoked potentials that are shared across all recordings due to reference contamination. This intervention prevents false positive connectivity results.

While it would be ideal to set the reference electrode *a priori* in electrically neutral tissue, this may not always be feasible due to the large quantity and variation in stimulation locations and signal spread across stimulation sites and conditions. In practice, only a subset of stimulation sites or conditions might be affected by reference contamination, which warrants addressing on a case-by-case basis.

Here, we focus on the issue of reference contamination by showcasing the occurrence of biased connectivity results in datasets collected at two different centers (Baylor College of Medicine, “Baylor”; Mayo Clinic, “Mayo”) with different recording and stimulation equipment but with the same issue: the presence of an apparent common evoked potential signal across many electrodes. We hypothesized that this issue was caused, in both cases, by the proximity of the reference electrodes to tissue displaying a strong response to stimulation.

Reference contamination can be reduced by *post hoc* re-referencing or by repeating the stimulation experiment using a different reference electrode. In this work, we tested both approaches. Specifically, at each center we repeated data collection after switching to a more neutral reference electrode, and focused on using the relationships between the two runs (contaminated vs. neutral reference) to validate the effectiveness of re-referencing. Our main assumption was that the runs collected with a neutral reference are closest to the ground truth. Following this assumption, we tested the efficacy of a data-driven re-referencing method in recovering that ground truth, based on three key predictions. First, the contaminated and neutral reference runs, which were independently acquired, should increase in similarity after re-referencing. Second, signals in the contaminated reference run should change by a greater degree after re-referencing compared with the neutral run. Finally, similarity metrics between all channels within a run should decrease with a neutral reference and after re-referencing, reflecting the natural variability of stimulation responses across the brain.

With this work, we aim to showcase different solutions to correct the issue of a contaminated reference in SPES experiments and offer a pipeline to address this issue in existing recordings. By comparing the results obtained using a neutral reference versus data-driven re-referencing scheme, we demonstrate that the re-referenced PEPs have similar features to the PEPs recorded using a neutral reference.

## Materials and Methods

### Subjects

Two human subjects provided informed consent to participate in this study while undergoing intracranial epilepsy monitoring with stereo-electroencephalography (sEEG). One subject (22-year-old male) was assessed at Baylor College of Medicine, Houston TX, and the other (18-year-old female) at Mayo Clinic, Rochester MN. Experimental procedures were conducted in accordance with the policies and principles outlined in the Declaration of Helsinki and were approved by the Institutional Review Board at Baylor College of Medicine (H-18112) and the Institutional Review Board of the Mayo Clinic (IRB 15-006530). SPES experiments were conducted while the patients were on anti-seizure medications and interictal epileptic activity was at its minimum. At Baylor, these experiments occurred one day after electrode implantation, while at Mayo, they occurred one day before electrode explantation.

### iEEG data collection

#### Baylor - Subject 1

The participant underwent surgical placement of 15 sEEG depth probes, with 185 total electrode contacts spanning various anatomical locations across left frontal and bilateral temporal regions. Ten probes had a 0.8 mm diameter with 8-16 electrode contacts of 2 mm length and 3.5 mm center-to-center distance (PMT Corporation, MN, USA); the 5 remaining probes had a 1.28 mm diameter with 9 recording contacts of 1.57 mm length and 5.0 mm center-to-center distance between contacts (AdTech Medical Instrument Corporation, WI, USA). Neural signals from all sEEG probes were recorded using a 256-channel Cerebus system (Blackrock Microsystems, UT, USA) at a 30 kHz sampling rate, with a 4^th^ order Butterworth high pass filter (0.3 Hz).

#### Mayo - Subject 2

The participant underwent surgical placement of 13 sEEG depth probes, with a total number of 207 electrode contacts spanning various anatomical locations across left frontal and bilateral temporal regions. All probes had a 0.8 mm diameter with 15-18 electrode contacts of 2 mm length and 3.5 mm center-to-center distance (DIXI medical, Marchaux-Chaudefontaine, France). Neural signals from all sEEG probes were recorded using a 256-channel g.HIamp system (g.tec, Schiedlberg, Austria) at a 4800 Hz sampling rate.

### Single pulse electrical stimulation

#### Baylor - Subject 1

We employed a monopolar cathodic SPES paradigm. Biphasic symmetric pulses with 5 mA amplitude, 180 μs pulse width, and a 100 μs interphase gap were delivered to an electrode contact in the right anterior hippocampus (stimulation seed) using a Blackrock CereStim R96 stimulator (Blackrock Microsystems, Utah, USA). Fifty single pulses were delivered (trials) with a variable inter-stimulation period ranging uniformly between 400 ms and 800 ms. We performed a total of 2 experimental runs using different reference and ground electrodes: run 1 employed two electrode contacts visually determined to be in white matter as the reference and ground, located 12.5 mm away from the stimulation site; run 2 employed two electrode contacts along one sEEG probe (not employed for recordings) placed in the subgaleal space (midline between the skin and the skull bone). Runs 1 and 2 are referred to from here on as having contaminated and neutral references, respectively. They were performed on the same day, spaced 1 hour apart.

After acquisition, signals were downsampled to 2kHz. Data from electrodes used as ground and reference were removed from analysis, along with 4 other electrode contacts that demonstrated poor signal quality.

#### Mayo - Subject 2

We employed a bipolar SPES paradigm. Biphasic symmetric pulses with 6 mA amplitude, 100 μs pulse width and no interphase gap were delivered to an electrode pair in the thalamus using a g.Estim PRO electrical stimulator (g.tec, Schiedlberg, Austria). Twelve single pulses were delivered with a variable inter-stimulation period ranging uniformly between 2.4 s and 5 s. We performed a total of 2 experimental runs with different reference electrodes (which also acted as the ground) visually determined to be in the white matter. The distances from each reference electrode to the center of the bipolar stimulation site were 43.2 mm (run 1) and 42.7 mm (run 2). As with the Baylor dataset, runs 1 and 2 are referred to from here on as having contaminated and neutral references, respectively. They were similarly performed on the same day, spaced one hour apart.

Signals were kept at the native 4800 Hz sampling rate, high pass filtered by forward-reverse filtering (2nd order Butterworth with cutoff frequency = 0.3 Hz), and had line noise attenuated by a spectrum interpolation technique (Mewett et al., 2001). Data from electrodes were removed if they displayed poor signal quality or consistent interictal activity, or if the electrodes acted as ground for either run. Additionally, 2 of the 12 trials were removed from the neutral reference run due to interictal activity.

### Theory and Calculation of Re-referencing

We implemented a data-driven re-referencing approach, termed CARLA, detailed in a recent publication (Huang et al., 2024). CARLA is an adjusted common average re-referencing technique that selects a subset of channels from the input dataset to use as an average reference. The average reference produced is an optimized tradeoff between reducing shared noise between all channels and minimizing bias from highly responsive or artifactual channels. Compared to common average and bipolar re-referencing, CARLA incurs less risk of distorting the evoked potential shape of individual channels. Briefly, all channels are ranked in order of increasing stimulation responsiveness by mean cross-trial covariance, and an increasing number of least responsive channels are iteratively taken to construct a common average. A test statistic based on Pearson correlation captures the anticorrelation between individual channels and the re-referenced subset at each iteration. Using too few channels for the common average increases anticorrelation due to overrepresented noise from each constituent, while using too many channels also increases anticorrelation due to the overinfluence of highly responsive ones. Thus, a peak in the test statistic detects the optimal largest subset that stops short of highly responsive channels.

In order to more robustly address reference contamination, we added an initial step before applying CARLA: all channels are first re-referenced to the median channel by responsiveness (mean cross-trial covariance). In case of a contaminated reference, this median channel’s waveform likely captures the reference contamination. Without this initial step, CARLA may at times stop short of capturing the reference contamination in its optimal subset by only including a small number of artifactually silent channels. These silent channels correspond to electrodes that are physically close to the same responsive tissue as the reference electrode and see their PEPs effectively “canceled out” by the reference signal.

### Analyses

#### Between-run difference

Statistical difference between experimental runs (contaminated vs. neutral reference) was quantified with the two-sample *t*-statistic between runs, calculated from the PEP waveforms recorded from each channel at each time point:

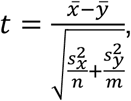

where 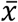 and 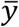 are the trial means for the contaminated and neutral reference runs, respectively, *s*_*x*_ and *s*_*y*_ are the standard deviations across trials, and *n* and *m* are the number of trials in each run. This resulted in a channels by time points *t*-statistic heatmap. Average difference post-stimulation was then summarized by averaging the absolute *t*-statistic values between 9 and 200 ms post-stimulation for each channel. We compared between-run differences before and after re-referencing with CARLA (contaminated vs. neutral, contaminated CARLA vs. neutral CARLA).

#### Cross-channel correlation

Cross-channel correlation heatmaps were created by averaging PEP waveforms across trials and calculating Spearman’s rho (ρ), from 9 to 200 ms post-stimulation, between all pairs of channels. This was done separately for each reference condition (contaminated, neutral, contaminated CARLA, neutral CARLA).

#### Latency detection

PEP components recorded intracranially are less established compared to CCEPs components recorded on the cortical surface (i.e., N1, N2, etc), especially given that the polarity of the deflections varies substantially depending on the cortical folding geometry surrounding the electrodes. Here, we focused on identifying the most prominent positive and negative peaks occurring in the time window spanning 9-100 ms after the stimulation pulse, without any assumptions regarding their association to CCEP components, and separately for each reference condition. To this end, we first averaged PEP waveforms across trials and applied temporal smoothing to each channel by subtracting the high-pass (>300 Hz) waveform. We then identified the largest occurring negative and positive peaks in the time window of interest with minimal prominence >20 μV, and measured their latencies with respect to the stimulation pulse. This resulted in a distribution of latencies across all channels for each reference condition. To assess the similarity in the peak latencies of PEPs across channels, two metrics of each distribution were computed: the percentage of latencies equal to the mode latency (mode size) and the entropy of the distribution.

The mode latency was identified for each distribution after first rounding latencies to the nearest millisecond integer. We then calculated the relative percentage of rounded latencies equal to the mode, out of all channels with that component identified (satisfying the peak prominence criterion).

To compute the entropy for each latency distribution, we first calculated the relative proportion, 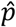, of latency in each 5 ms bin (5-100 ms). The entropy was then calculated as:

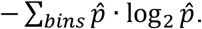

This operation was performed on bootstrapped resamples (n = 10,000) across channels to estimate mean and standard error.

#### Response durations

We applied the canonical response parameterization (CRP) method to measure the response durations of PEPs (9-1000 ms post-stimulation) in subject 2, separately for each reference condition (Miller et al., 2023).

The CRP method generates a response duration for each PEP that is based on voltage reliability across stimulation trials and indifferent to response shape. Response durations were only kept for PEPs with a significant response, as quantified by mean parameterized trial projection weights significantly different from 0 on average (one sample *t-*test, *p* < 0.05). We quantified mode size and entropy of the response duration distribution as for peak latency above. In this case, response durations were rounded to the nearest 10 milliseconds before computing the mode, and the relative percentage equal to the mode was computed out of all channels with significant response durations identified. Entropy was calculated on bins of 50 ms width (0-1000 ms).

## Results

We collected iEEG data from two subjects across two institutions with major differences in recording equipment and SPES experimental protocol (Baylor: subject 1, Mayo: subject 2). In each subject, one stimulation site was chosen for SPES and data were collected for two independent experimental runs in which the hardware reference electrode differed. The first run in each case exhibited a consistent signal across most PEPs suspected to be due to reference contamination, while the second run used a more neutral reference electrode. Our objective is to present several intuitive signal-derived metrics that could be used to detect the presence of a reference contamination, and to quantify the effectiveness of switching the hardware reference and/or *post hoc* re-referencing as mitigation strategies.

### Re-referencing increases PEP similarity between experimental runs

PEPs in both experimental runs are shown for each subject before and after re-referencing. (Figure 1, top). In run 1 (contaminated reference) for subject 1, a prominent negative deflection is observed across most channels, centered at 50 ms post-stimulation. This deflection is not present in run 2 (neutral reference) or in either run after re-referencing. Similarly in subject 2, the most prominent feature shared across channels in run 1 is a positive deflection centered at 120 ms post-stimulation. This feature also disappears after switching to a neutral reference in run 2 or after re-referencing, though a substantial amount of periodic noise is still present in run 2 before re-referencing.

**Figure 1.**
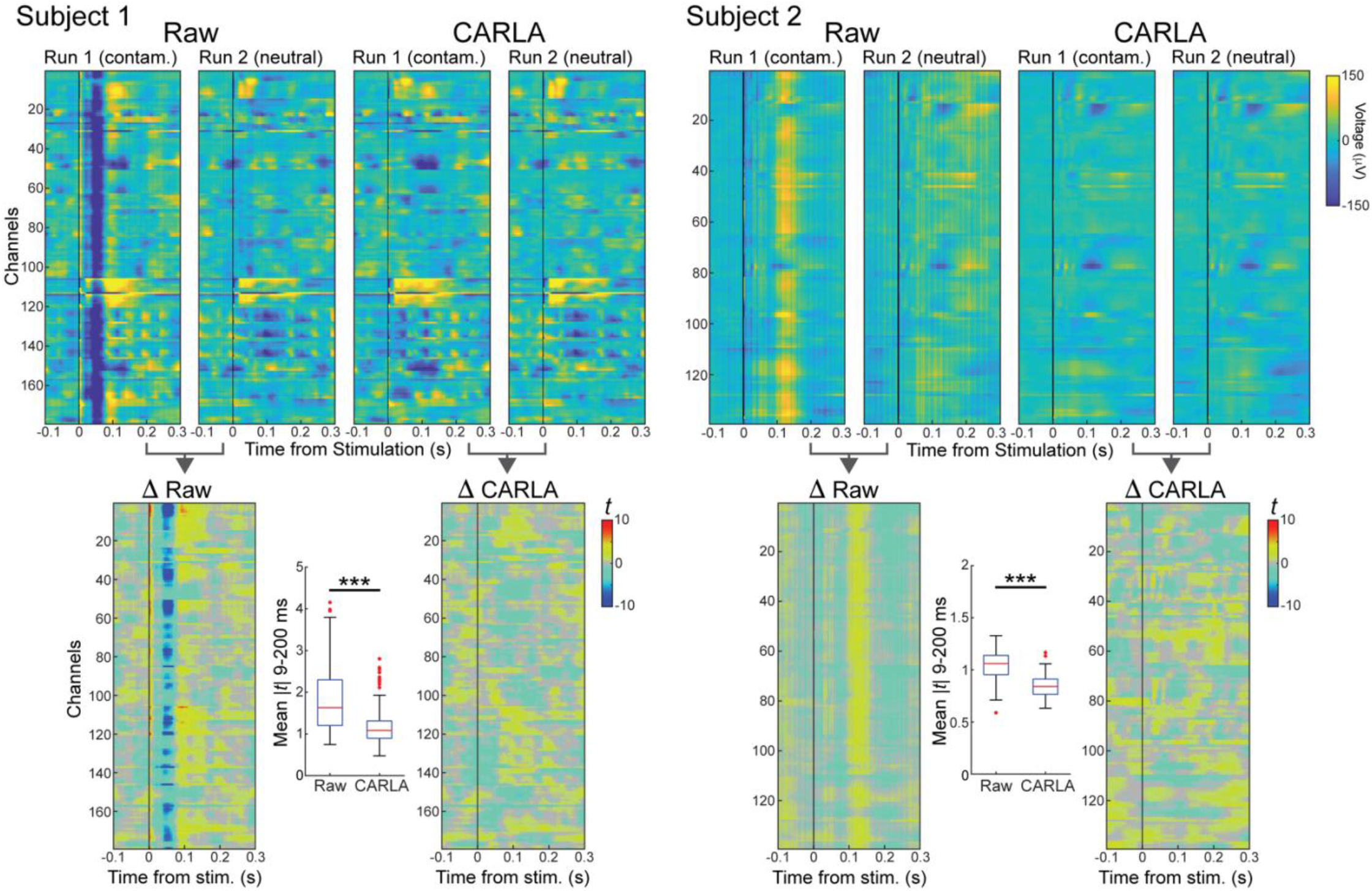
Examples of reference contamination and increased between-run similarity after re-referencing. **Top:** Mean pulse-evoked potentials (PEP) measured for all recording channels (y-axis): negative deflections are associated with blue colors, positive with yellow colors and the stimulation pulse time is indicated by a vertical black line (time 0 on the x-axis). Columns show mean PEPs obtained from recordings with a contaminated reference (run 1) and with a neutral reference (run 2), before (Raw) and after (CARLA) re-referencing. Subject 1: dataset from Baylor; Subject 2: dataset from Mayo. The presence of a common deflection across almost all channels (negative for Subject 1, positive for Subject 2) is indicative of a contaminated reference and so it is not present when using a neutral reference or after re-referencing. **Bottom:** Per-time point *t*-statistics quantify PEP differences between contaminated and neutral reference runs, before and after re-referencing. Boxplots show mean |*t*| between 9-200 ms for each channel. The difference between runs is large in the raw data; after re-referencing with CARLA the differences are significantly reduced, indicating that the influence of the contaminated reference was effectively removed and mean PEPs became similar. *** indicates significant decrease in median (mean |*t*|) after re-referencing (Wilcoxon signed-rank test, *p* < 0.001).

PEPs are visually more similar between experimental runs in each subject after re-referencing than before. We quantified differences between runs on a per-channel, per-time point basis using the two-sample *t*-statistic (Figure 1, bottom). Larger magnitude *t*-statistic, positive or negative, indicates a greater difference between runs. The average difference post-stimulation was computed for each channel by averaging the absolute *t*-statistic value between 9 and 200 ms post-stimulation (Figure 2 boxplots). In both subjects, the average difference decreased significantly across channels after data were re-referenced. This convergence in PEP waveforms between experimental runs represents a successful correction of reference contamination by re-referencing.

**Figure 2.**
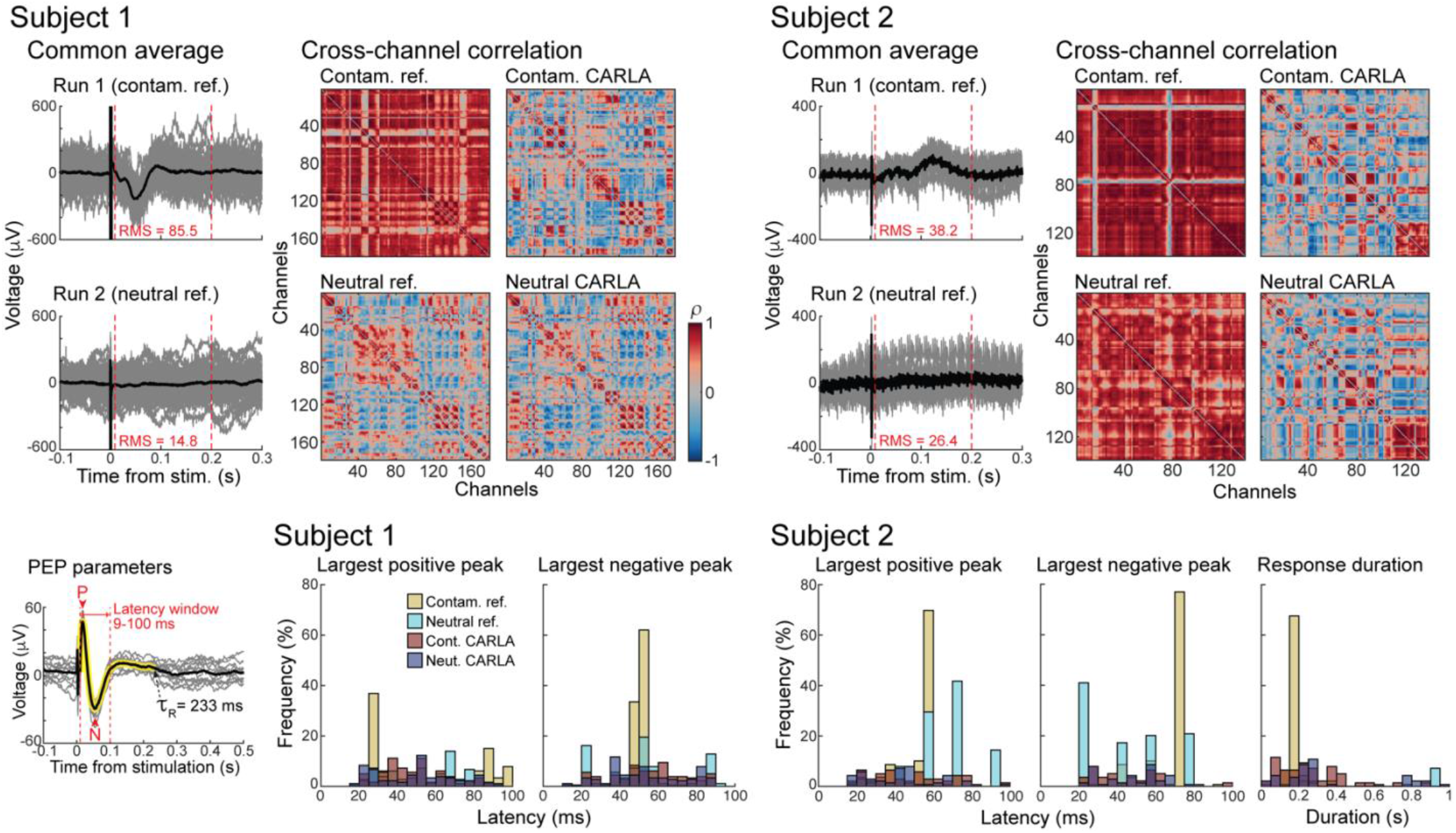
Cross-channel similarity is decreased with neutral reference, re-referencing, or both together. **Top:** Adjusted common averages determined by CARLA for contaminated and neutral reference runs (individual trials in gray, mean across trials in black). Root mean square voltage is calculated between 9-200 ms for the mean. Spearman’s rho (ρ) of PEPs (9-200 ms) between all pairs of channels for each run, before and after re-referencing with CARLA. **Bottom:** Distribution across channels of PEP parameters (latency of positive and negative peaks, response duration, illustrated in inset) in each run before and after re-referencing. Onset latencies of largest positive and negative peaks <100 ms (P and N in inset) were calculated for both subjects, and response durations estimated using the CRP method (*τ*_*R*_) were calculated for subject 2 only.

### Smaller adjusted common average with neutral reference than contaminated reference

In our modified implementation of CARLA re-referencing, an adjusted common average is constructed for each dataset using all channels that are relatively non-responsive, relative to the PEP waveform in the median channel. Therefore, any deflections in the adjusted common average should reflect the reference contamination shared across most channels rather than true PEPs. In both subjects, the adjusted common average is of lower root mean square amplitude, post-stimulation, for the neutral reference data than the contaminated reference data (Figure 2, top). This indicates a relatively smaller change necessary to correct PEPs when recording with a neutral reference. As expected, the deflections in the adjusted common averages for the contaminated reference runs match the prominent deflections seen across channels in Figure 1.

### Cross-channel correlation decreases after correcting for reference contamination

Spearman’s rho was calculated for all pairs of channels, 9-200 ms post-stimulation (Figure 2, heatmaps). This yielded 15,931 and 9,591 channel pairs for subjects 1 and 2, respectively. In subject 1, 96.6% of all channel pairs showed significant positive or negative correlation (*p* < 0.01, uncorrected) in run 1 (contaminated reference), with a mean correlation of rho = 0.67. Correlations were reduced by switching to a neutral reference (76.8% significant, mean rho = 0.08), by re-referencing alone (80.4% significant, mean rho = 0.01), and by performing both (78.6% significant, mean rho = 0.04).

In subject 2, 97.6% of all channel pairs showed significant correlation in run 1, with mean rho = 0.78. Switching to a neutral reference reduced the mean correlation (mean rho = 0.68) but did not reduce the overall fraction of significant pairs (98.5%). Both the fraction of significant pairs and mean correlation were reduced by re-referencing alone (87.0% significant, mean rho = 0.01), and by re-referencing after switching to a neutral reference (89.3% significant, mean rho = 0.03).

A high average correlation across pairs of channels reflected spurious global similarity that was driven by the presence of a common signal, seen in the adjusted common average for each run. In subject 1, both hardware and software solutions were similarly effective at reducing the correlative structure between channels. However, re-referencing was the more effective solution in subject 2. This was because of a residual correlation from high-amplitude common noise (e.g., 60 Hz line noise) specific to that recording environment, and this noise was independent of the common signal from the reference.

### PEPs vary more across channels after correcting for reference contamination

PEP Parameters were calculated for all channels in each referencing condition, for each subject (Figure 2, bottom). These consisted of the latencies of the most prominent positive and negative peaks (9-100 ms), and the overall response duration as quantified by the CRP method (9-1000 ms, subject 2 only). Detailed descriptive statistics of the PEP parameter distributions are listed in table S1.

Peak latencies measured in the presence of a contaminated reference appear highly concentrated around one or few values, which match the timing of peaks in the common averages. Two metrics that reliably capture this artifactual similarity in peak latencies (and thus potential reference contamination) are the percentage of values equal to the mode (mode size) and the entropy. The contaminated reference PEPs produced positive and negative peak latencies with a high percentage of all values equal to the mode (range: 21.7-77.0%). The mode size decreased drastically by switching to a neutral reference (6.6-41.7%) or by re-referencing (5.7-8.8%). Entropy quantifies the randomness of each distribution as a whole, indifferent to the exact number or separation of modes. Lower entropy indicates that peak latencies are more concentrated around one or more central values, while higher entropy indicates that peak latencies are more spread out. In all cases, entropy was lowest for the contaminated reference condition (1.13-2.31), and increased for neutral reference (1.93-3.41) and re-referencing (3.07-3.63) conditions. Both mode size and entropy reveal the high degree of PEP similarity across channels incurred by reference contamination. Combining a neutral reference and re-referencing produced results similar to re-referencing alone, and did not further improve the spread of PEP parameters (mode size: 5.3-9.8%, entropy: 2.93-3.57).

Switching to a neutral reference was as effective as re-referencing on decreasing mode size and increasing entropy for latencies in subject 1. However, re-referencing was the superior solution in subject 2. This was due to the high amplitude of common noise in subject 2, attenuated by re-referencing but not by switching reference electrodes alone. In fact, the spacing between the individual modes before re-referencing matches the period of line noise (16.7 ms). This difference in subject 2 is consistent with the pattern observed across correlation heatmaps.

With longer inter-stimulation intervals in subject 2, we also calculated response durations of PEPs using the canonical response parameterization (CRP) method. The mean response durations converged between the contaminated and neutral reference runs after re-referencing each with CARLA (from 171 ms and 667 ms to 273 ms and 350 ms). The mode size and entropy of response durations showed similar patterns across reference conditions as peak latencies: mode size decreased (from 60.3%) with neutral reference (29.4%), re-referencing (7.7%) or both (14.5%); and entropy increased (from 1.05) with neutral reference (1.53), re-referencing (3.34), or both (2.77).

## Discussion

In this work, we bring attention to a type of referencing issue that can be present in PEP data. When the hardware reference electrode is contaminated with a time-locked signal, that signal propagates to all other channels and leads to erroneous conclusions about timing and connectivity. We presented ways to identify the reference contamination and demonstrated that the issue can be resolved by hardware and software solutions.

In the ideal scenario, all data are recorded relative to a neutral reference electrode in one experimental session. However, this is difficult to achieve consistently in practice. We found that simply choosing a reference electrode far from the stimulation site is not sufficient, because reference contamination can arise from proximity to responsive tissue, which can occur at distant locations (Paulk et al., 2022). When all stimulation sites are unilateral, selecting a reference electrode in the opposite hemisphere may be effective. Or, as seen for subject 1, selecting an electrode outside of brain tissue – in this case in the subgaleal space – is also a viable solution. However, when a neutral reference electrode is not available, it becomes crucial to identify reference contamination and employ mitigation strategies. If all recording channels show very similar response dynamics, reference contamination should be suspected. We recommend incorporating a simple step to the experimental pipeline, wherein the outgoing trial-averaged PEPs from each stimulation site are plotted for all channels immediately after data acquisition. This should be done even if analysis only requires single recording channels. Reference contamination may present more subtly than shown here, with lower amplitude, shorter duration, or mixed with true PEPs. The reference signal may also be “canceled out” at a subset of electrodes in similar electrophysiological environments as the reference.

Reference contamination can be addressed in several ways. The simplest approach is to recollect data with a different hardware reference electrode, as in this study, though this is often impractical due to the unique time constraints of human iEEG. *Post hoc* re-referencing is therefore a practical alternative. One common version is bipolar re-referencing, which is computationally efficient but can attenuate or distort true PEPs that are spatially adjacent (Arnulfo et al., 2015; Shirhatti et al., 2016). Instead, this manuscript highlights an optimally adjusted common average re-referencing approach, CARLA, which eliminates globally shared reference contamination with minimal impact on the underlying PEP waveforms.

We evaluated the effectiveness of our solutions using several metrics based on our initial predictions. First, the per-sample *t*-statistic heatmaps showed that the two experimental runs agreed more with each other after both were independently re-referenced. Second, the adjusted common averages were of lower average amplitude and did not show prominent deflections when the data were collected with a neutral reference. Third, PEPs showed less overall correlation across all channels, and PEP parameters demonstrated greater natural variability across the brain when using a neutral reference, CARLA re-referencing, or both. These solutions were effective regardless of the stimulation paradigm – monopolar cathodic at Baylor and bipolar at Mayo – highlighting their versatility for diverse clinical and research settings. Across settings, PEP latencies and response durations are used to interpret interareal connectivity, making it crucial to ensure that they accurately reflect physiological phenomena. Lastly, we note that re-referencing provides two additional advantages with respect to switching the reference electrode: it can mitigate a contaminated reference issue on existing data and it removes other types of unwanted features that are common across channels, such as residual line noise.

## Conclusion

Our results demonstrate the importance of selecting a neutral reference location to avoid biased connectivity results. The high variability between iEEG recording methods, amplifiers and in-house software packages makes quality checks challenging. Data can be collected with grids or depth electrodes and stimulation parameters can vary in amplitude, pulse duration, inter-pulse interval and mono- or bipolar configurations. This manuscript provides a clear set of quality checks for PEP data and an open software solution to fix the issue. We validated these methods on data collected with different equipment at two different institutions. In both settings, we demonstrate how careful data-driven re-referencing tools can be used to remove the influence of the reference electrode and obtain unbiased connectivity results from PEPs.

## Funding

This work was supported by the National Institutes of Health [R01-MH122258 and R01-EY035533 to D. H., R01-MH127006 to K. R. B., T32-GM145408 to the Mayo Clinic MSTP, U01-NS128612 to K. J. M., U01-NS103549 to S. A. S] and by the McNair Foundation (S. A. S.).

## CRediT

**Harvey Huang:** Conceptualization, Formal Analysis, Methodology, Software, Validation, Visualization, Writing – Original Draft, Writing – Review & Editing. **Joshua Adkinson:** Investigation, Methodology, Writing – Review & Editing. **Michael A. Jensen:** Methodology, Validation, Writing – Review & Editing. **Mohammed Hasen:** Investigation, Resources, Writing – Review & Editing. **Isabel A. Danstrom:** Investigation, Writing – Review & Editing. **Kelly R. Bijanki:** Conceptualization, Funding Acquisition, Writing – Review & Editing. **Nicholas M. Gregg:** Investigation, Supervision, Writing – Review & Editing. **Kai Miller:** Conceptualization, Funding Acquisition, Methodology, Resources, Software, Writing – Review & Editing. **Sameer A. Sheth:** Conceptualization, Funding Acquisition, Resources, Writing – Review & Editing. **Dora Hermes:** Conceptualization, Data Curation, Formal Analysis, Funding Acquisition, Investigation, Methodology, Software, Supervision, Writing – Original Draft, Writing – Review & Editing. **Eleonora Bartoli:** Conceptualization, Data Curation, Formal Analysis, Methodology, Supervision, Validation, Writing – Original Draft, Writing – Review & Editing.

## Declaration of competing interest

S. A. S. is a consultant for Boston Scientific, Neuropace, Koh Young, Zimmer Biomet, Varian Medical, and Sensoria Therapeutics and co-founder of Motif Neurotech. K. R. B. has one issued patent and one patent pending, both unrelated to the current work (US 63/592,453, and US 11,241,575). N. M. G. has consulted for NeuroOne, Inc. (funds to Mayo Clinic). The authors declare no other competing interests.

## Supplementary Information

**Table S1.** PEP Parameters across reference conditions.

|  | Positive Peak |  |  | Negative Peak |  |  | Response Duration |  |  |
| --- | --- | --- | --- | --- | --- | --- | --- | --- | --- |
| | Mean ± SD<br>(ms) | Mode<br>Size (%) | Entropy ±<br>SE | Mean ± SD<br>(ms) | Mode<br>Size (%) | Entropy ±<br>SE | Mean ± SD<br>(ms) | Mode<br>Size (%) | Entropy ±<br>SE |
| **Subject 1** |  |  |  |  |  |  |  |  |  |
| **Contam.** | 54.2 ± 30.2 | 21.7 | 2.31 ± 0.11 | 50.7 ± 3.4 | 57.0 | 1.20 ± 0.08 | | | |
| **Neutral** | 54.7 ± 18.6 | 6.6 | 3.41 ± 0.09 | 51.0 ± 21.8 | 12.8 | 3.15 ± 0.11 | | | |
| **Contam.**<br>**CARLA** | 49.5 ± 16.9 | 5.8 | 3.44 ± 0.09 | 54.8 ± 19.3 | 5.7 | 3.63 ± 0.07 | | | |
| **Neutral**<br>**CARLA** | 50.4 ± 20.3 | 7.3 | 3.55 ± 0.10 | 53.4 ± 20.1 | 5.3 | 3.57 ± 0.09 | | | |
| **Subject 2** |  |  |  |  |  |  |  |  |  |
| **Contam.** | 51.4 ± 10.9 | 69.8 | 1.55 ± 0.15 | 68.1 ± 13.4 | 77.0 | 1.13 ± 0.12 | 171 ± 44 | 60.3 | 1.05 ± 0.16 |
| **Neutral** | 65.7 ± 17.1 | 41.7 | 2.08 ± 0.11 | 45.1 ± 19.9 | 41.0 | 1.93 ± 0.06 | 667 ± 341 | 29.4 | 1.53 ± 0.33 |
| **Contam.**<br>**CARLA** | 49.4 ± 20.7 | 5.8 | 3.50 ± 0.11 | 44.4 ± 17.6 | 8.8 | 3.07 ± 0.11 | 273 ± 193 | 7.7 | 3.34 ± 0.11 |
| **Neutral**<br>**CARLA** | 43.4 ± 19.2 | 6.8 | 3.20 ± 0.15 | 48.2 ± 16.1 | 9.8 | 2.93 ± 0.17 | 350 ± 283 | 14.5 | 2.77 ± 0.15 |

